# Multiple locomotion gaits in the mole cricket

**DOI:** 10.1101/2025.01.19.633773

**Authors:** Omer Yuval, Avi Amir, Amir Ayali

**Author notes:** These authors contributed equally (Omer Yuval, Avi Amir). Author contribution: Avi Amir, Omer Yuval and Amir Ayali conceived the study and planned the experiments. Avi Amir collected and maintained the animals, designed, developed, and built the experimental setup, recorded the animals, annotated the videos, and established a database of mole cricket locomotion. Omer Yuval wrote the original draft, analyzed the experimental data, and developed the biomechanical model and the reinforcement learning algorithm for simulating insect locomotion. Amir Ayali secured the required funding and supervised the work. All authors reviewed the manuscript. The authors have no competing interests related to this study. Funding sources: This research was funded by the Isreal Science Foundation ISF, grant no. 391/24 to Ayali A.

## Abstract

Insects are exceptionally robust walkers. Although different species exhibit distinct anatomical and functional specializations, they are also highly adaptive within these constraints. How such adaptations enable insects to efficiently navigate diverse environments and perform mechanical tasks remains far from fully explored. The mole cricket, which dwells underground, is one of the least studied insects, largely due to its cryptic lifestyle. It excels at digging tunnels and exhibits extreme morphological adaptations, particularly its exceptional fossorial forelegs. Its versatile locomotion, above and below ground, makes the mole cricket an attractive model system for studying the biomechanics of insect movement. Here we provide the first quantitative characterization of mole cricket locomotion. Using a tunnel-like arena, we recorded freely-moving insects and analyzed their various locomotion gaits. We identified and described three main modes of locomotion, including a backward-bound gait that has not previously been reported in any insect. To test specific hypotheses regarding form-function relationships and the generation of thrust, we integrated biomechanical modeling and deep reinforcement learning to simulate the observed gaits. Our work opens several future directions, from exploring context-dependent gait transitions to bio-inspired technological innovations.

## 1 Introduction

The ability of legged animals to propel their body in a specific direction and at a certain speed is closely related to their body plan. Locomotion is constrained by body morphology, particularly the number of limbs, their size and segmentation, and the degree of freedom of the joints that connect them. Within these constraints, insects exhibit a great variation across species, manifested in the geometry and biomechanics of their body parts [1, 2]. Interestingly, the evolution of such anatomical variation is often coupled with the evolution of specialized neuromuscular control mechanisms, which together enable animals to move efficiently and survive in their natural habitats. These co-evolution processes have given rise to different locomotion gaits that may be advantageous in specific environments (e.g., in the air, in water, on surfaces, or underground) and contexts (e.g., during exploration, feeding, mate finding, or under stress conditions).

Insects possess a three-part body comprising a head, thorax, and abdomen, with six legs connected to the thorax. The segments of each leg are the coxa, femur, tibia, and tarsus, which are connected via joints and controlled by striated muscles [2, 3]. Stepping behaviors in insects, as in all legged animals, are typically described as a sequence of swings and stances, in which the legs touch the ground only during the stance phase [4]. The common forward walking via the alternating tripod gait is well documented and has been studied in many swift-moving insects [5, 6], such as cockroaches [7], flies [8], ants [9], and crickets [4]. In this gait, each triplet of non-adjacent legs typically moves in-phase and touches the ground at the same time, while being in anti-phase with the other triplet. However, measurements have shown some variation in inter-leg phases across species [4, 6, 10, 11].

The European mole cricket (*Gryllotalpa sp.*) is a 5 cm-long nocturnal insect that dwells underground and specializes in digging tunnels (Fig. 1A) [12–14]. It feeds on worms, small arthropods, and vegetable roots [15] and is considered an agricultural pest [16]. Its notable morphological adaptations include a pair of fossorial forelegs (Fig. 1A) and an enlarged pronotum — a shield-like chitinous exoskeleton that protects its head and thorax [17]. How these morphological adaptations affect the the insect’s locomotion, its ability to generate thrust, and nagotiate its unique environment is far from fully explored [14, 18–20]. Moreover, Kidd [15] has already noted the exceptional ability of the mole cricket to move backward, noting that “in order to prevent the necessity of its excavating a track so wide as to admit of the body being turned round in case of a desire to retreat, it is endued with the power of moving as easily in a retrograde as in a progressive direction”. There is relatively scarce evidence in the literature for back-ward locomotion in insects [21–23]. In these few documented cases inter-leg phases were shown to be more irregular compared to in forward walking [23], suggesting that most insects prefer to turn around rather than walk backwards. The mole cricket’s extreme morphological adaptations and its asymmetric and unique body plan thus raises questions regarding the co-evolution of body geometry and mechanics, and the locomotor gaits and control mechanisms used to navigate efficiently within the constraints of its natural habitat. It presents an ideal model for uncovering generalized form-function principles in insects.

**Figure 1.**
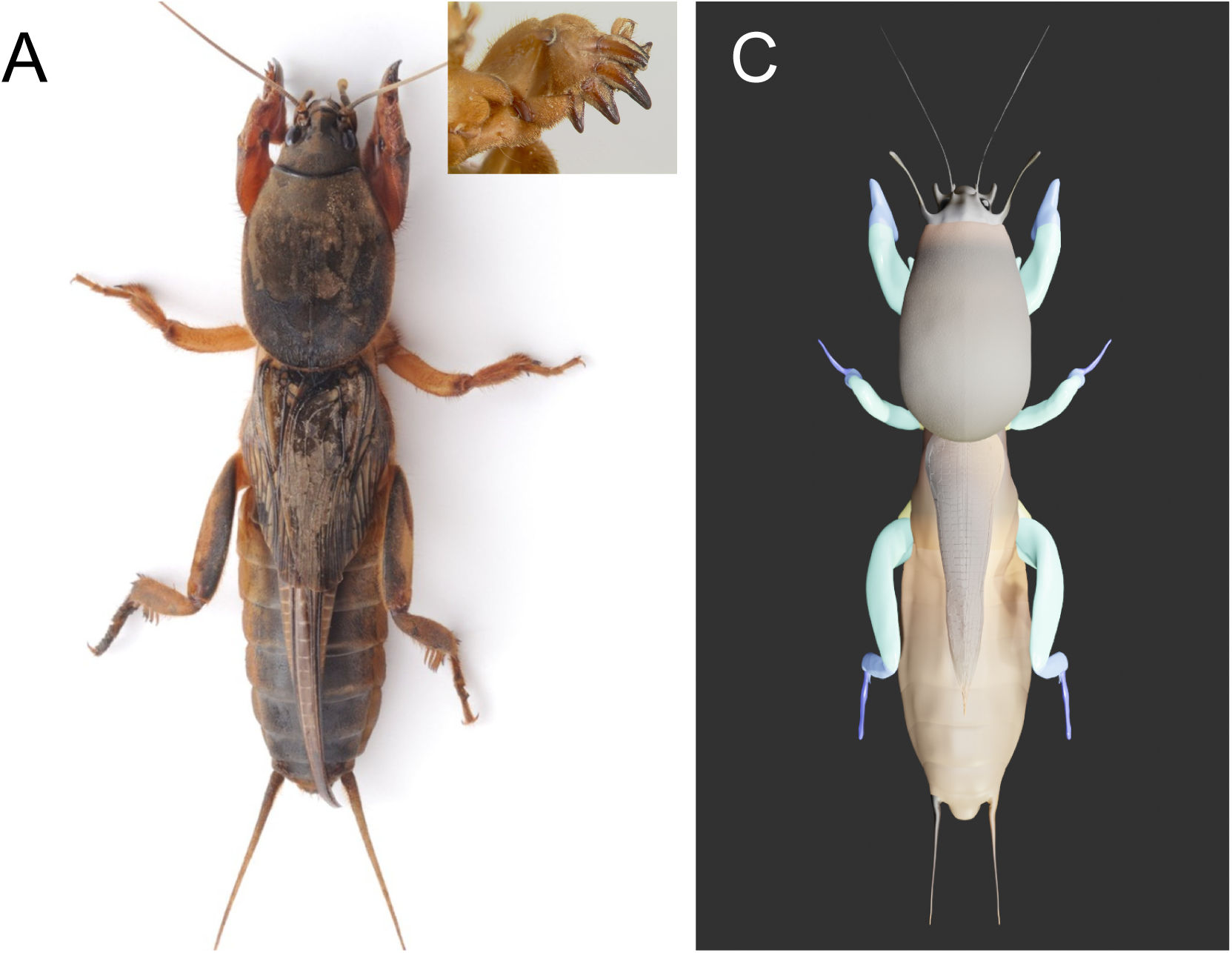
Studying mole cricket locomotion using behavioral assays and mechanical modeling. A. An adult mole cricket (*Gryllotalpa sp.*). B. The arena used to study its locomotion. Animals walk along the track, which has acclimatization chambers at either end. C. A 3D model of the mole cricket.

In this work we monitored and recorded the locomotion of male and female mole crickets in a tunnel-like environment (Fig. 1B). Insects were free to move within the tunnel section without additional constraints (such as tethering or a treadmill) [24]. Analysis of the recordings revealed three distinct walking gaits, including the recently described backward-bound gait [25] that has not been reported for any other insect to date. By tracking body and leg positions in the animal’s intrinsic coordinate system, we computed the speed of the whole body, stepping frequency, inter-leg phases, and leg trajectories. This allowed us to create the first corpus and quantitative characterization of mole cricket walking behaviors. We compared forward and backward double-tripod walking in order to examine the role of longitudinal body asymmetries and gait-specific kinematic parameters in the insect’s ability to propel itself forward or backward. We also demonstrated that the backward-bound gait is on average faster than the backward double-tripod walking, suggesting that it has evolved as an escape mechanism. Finally, in order to link specific kinematic parameters to the ability of the animal to move itself forward or backward in each gait, we developed a geometrically-accurate model of the mole cricket (Fig. 1C) and used mechanical modeling combined with deep reinforcement learning to simulate the observed gaits. This allowed us to test the effects of gait modifications that cannot be tested in vivo, and to make predictions regarding the role of specific legs and kinematic parameters in the ability of the animal to generate thrust.

## 2 Results

### Mole crickets use an alternating tripod gait to walk both forwards and backwards

To characterize the locomotory patterns used by the mole cricket, we recorded freely-moving adult males and females in a tunnel-like arena (Fig. 1B). To extract the spatio-temporal features of the mole cricket’s locomotion, we used machine learning to track the position of the base and tip of each leg, the head-thorax junction, and the thorax-abdomen junction (Fig. 2A and see Methods). The mole cricket, though bulky in appearance, was found to be remarkably agile when walking. As previously reported [14], despite its unwieldy fossorial forelegs, we found that the mole cricket employs the common double-tripod gait to move forward (Fig. 2B, F). Interestingly, the insect also uses a double-tripod pattern when walking backwards (Fig. 2C, G). In contrast to other insects that walk backwards, the backward gait in the mole cricket remains stable for many cycles (up to 9 cycles in our setup, restricted by the tunnel length). As detailed below, backward walking is not simply forward double-tripod gait “played in reverse”; and the two gaits differ in various kinematic features (beyond the principal walking direction)

**Figure 2.**
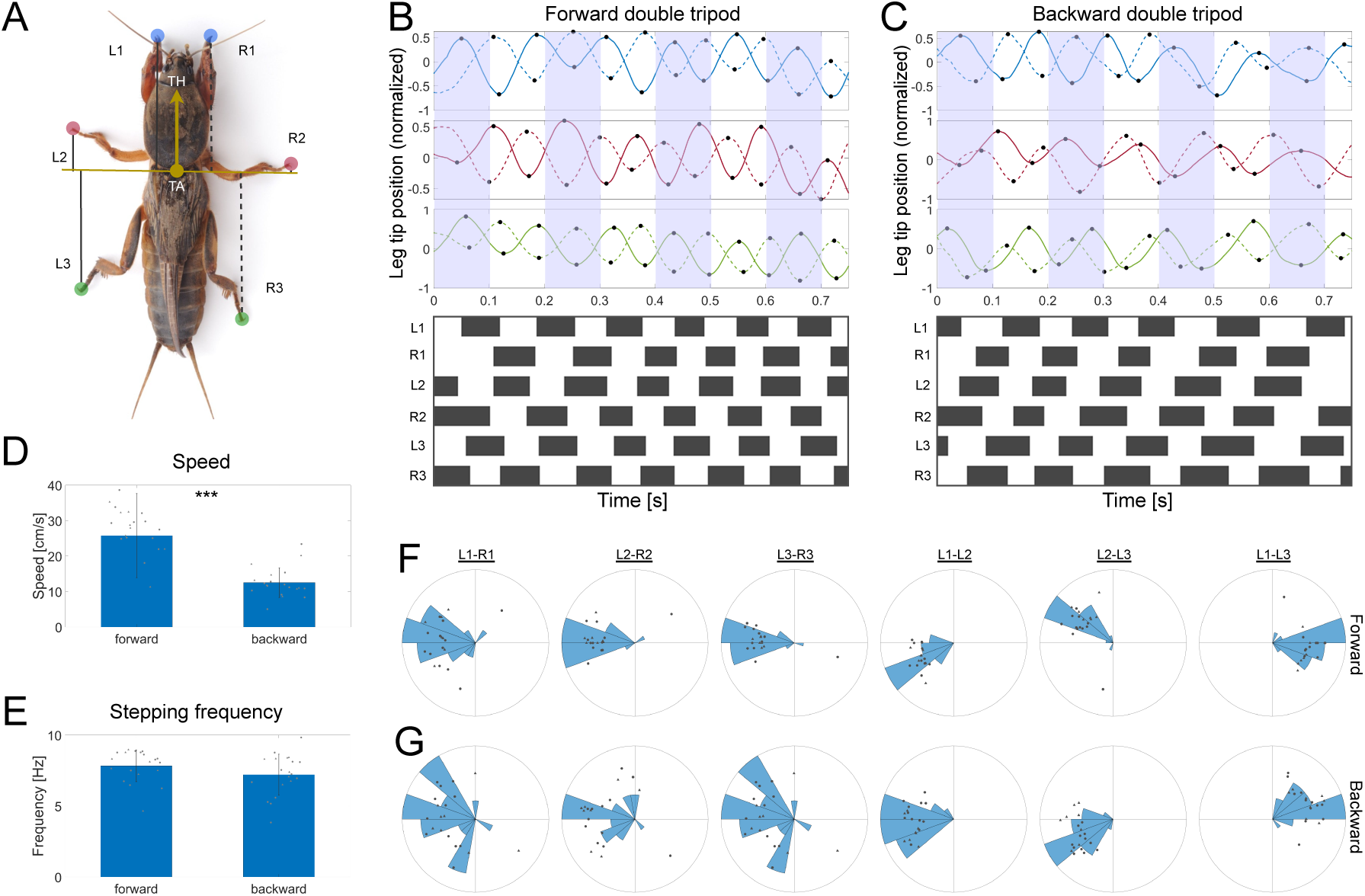
Mole crickets use an alternating tripod pattern to walk forwards and backwards. A. A diagram of the labeled body points, showing the thorax-head junction (TH), thorax-abdomen junction (TA), and leg tips (forelegs in blue, middle legs in red, and hind legs in green). The black lines show the distance, *d_i_*, of each leg tip from the TA along the main body axis (yellow arrow). B-C. Example of leg tip distances over time, *d_i_*(*t*), in forward (B) and backward (C) locomotion, showing the forelegs (top, blue), middle legs (middle, red), and hindlegs (bottom, green), and corresponding swing stance diagrams (bottom, black bars show stances; white bars show swings; see Methods). D-E. Mean whole body speed (D) and stepping frequency (E). F-G. Histograms of inter-leg phases in forward (F) and backward (G) locomotion (see Methods). Dots show mean inter-leg phases for single clips. Statistics were calculated using the two-sample t-test. \*\*\**p <* 0.0005. n=20 clips of different animals per gait, including at least 8 females (circles) and 8 males (triangles). Bars show the mean value across animals; error bars show the standard deviation.

To compare the insect’s ability to generate thrust between the two gaits, we measured whole-body speed (see Methods) and found that forward walking was on average, significantly faster than backward walking (Fig. 2D). We then asked whether the reduced speed in backward walking could be explained by a smaller number of step cycles per unit time. Surprisingly, we found that stepping frequency did not differ significantly between the two gaits (7.8 ± 1.2Hz vs 7.2 ± 1.4Hz; Fig. 2E). This suggests that the difference in speed stems either from other leg kinematic properties (not reflected in stepping frequency), or from morphological asymmetries along the main body axis (i.e., anterior-posterior differences in leg’s and overall body shape).

### Middle legs trajectory orientation and span can explain speed reduction in backward locomotion

Given that forward walking is faster than backward walking despite similar stepping frequencies (Fig. 2D-E), we asked which leg kinematic properties might be associated with whole-body speed. To assess the contribution of the different legs to forward and backward speed, we examined the shape and orientation of their leg tip trajectories (Fig. 3A-B and see Methods). We observed striking differences, which were most pronounced in the middle legs (Fig. 3A-B). Furthermore, to approximate the amount of force applied to the ground during each step cycle, we measured the area of the convex contour of each leg trajectory (Fig. 3A-B, gray contours; see Methods), and found that during forward locomotion, the middle legs covered, on average, a larger area (Fig. S1A).

**Figure 3.**
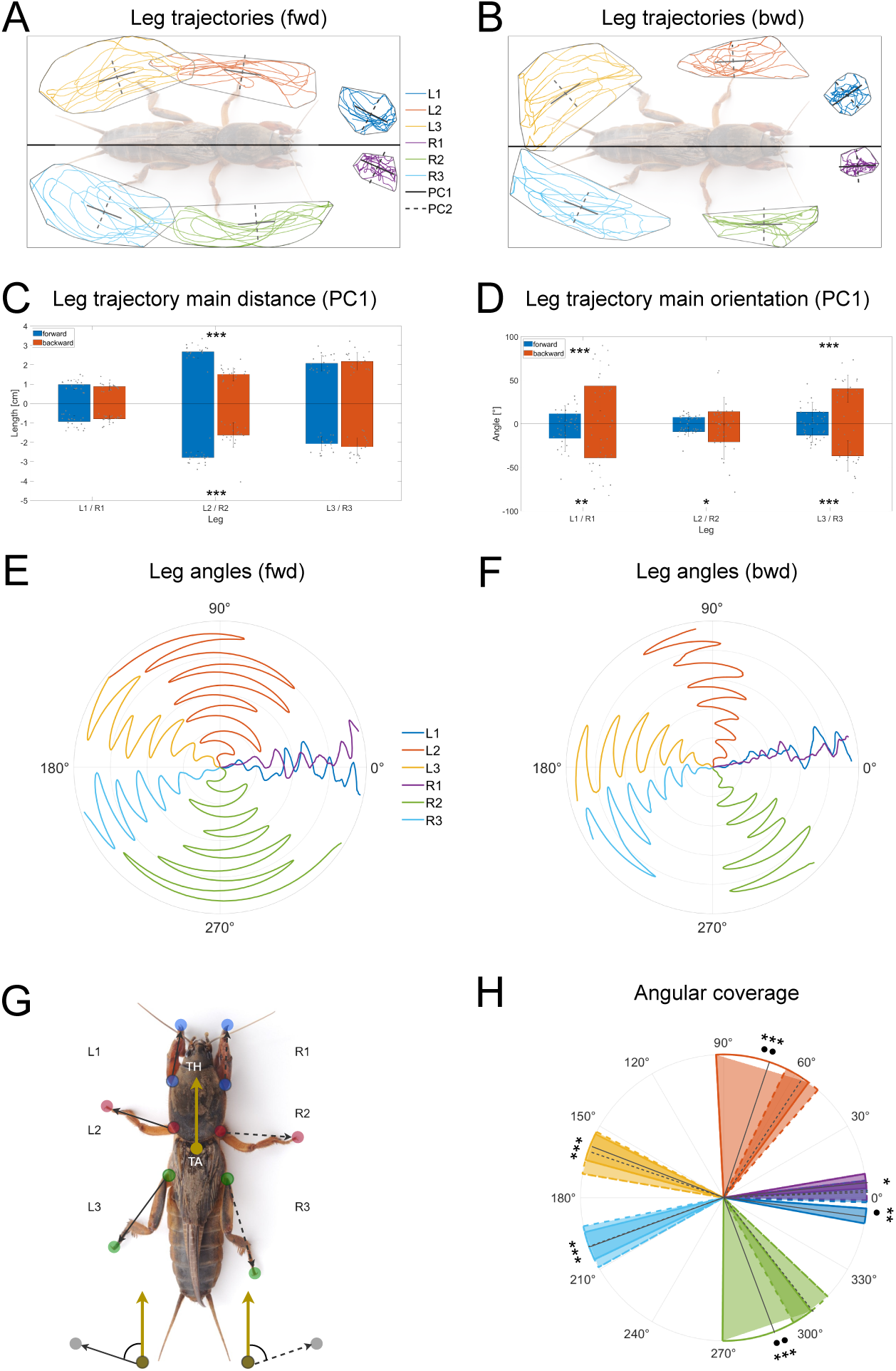
Spatial characterization of forward and backward alternating tripod locomotion. A-B. Leg trajectories of forward (A) and backward (B) locomotion (see Methods). The gray contour lines show the minimal convex contour of each leg trajectory. The black solid and dashed lines show the primary (PC1) and secondary (PC2) directions of each leg trajectory, respectively (see Methods). C. Leg trajectory distance along the primary direction (PC1; see Methods). D. Leg trajectory primary orientation. E-F. Examples of leg angles in forward (E) and backward (F) locomotion (see Methods). Time is shown in the radial coordinate. G. A diagram showing leg angles, defined as the angle between the vector connecting the base (colored circles) to the tip (colored circles with borders) of the leg (black arrows) and the primary body axis (yellow arrow). H. The mean angular sector covered by each leg in forward (solid lines) and backward (dotted lines) locomotion. The circular sectors show the average standard deviation of leg angles across all animals, and their center shows the average leg orientation (black solid lines for forward and black dashed lines for backward). Statistics were calculated using the two-sample t-test. \**p <* 0.05, \*\**p <* 0.005, \*\*\**p <* 0.0005. Bars show the mean value across animals, and error bars show the standard deviation. In I, statistics for the mean leg angles were calculated using Kuiper’s test. ●*p <* 0.05, ●●*p <* 0.005. Stars show statistically significant difference between the standard deviations of leg angles. n=20 clips of different animals per gait, incorporating at least 8 females (circles) and 8 males (triangles).

To estimate the contribution of stance properties to whole-body speed, we used principal component analysis (PCA) to decompose each leg trajectory into its primary (PC1) and secondary (PC2) components (see solid and dashed lines in Fig. 3A-B). PC1 was used to approximate step stride angle and PC2 was used to approximate side motion. We found that middle-leg trajectories traced larger distances along the PC1 direction in forward versus back-ward locomotion, while there was no significant difference in the forelegs and hindlegs (Fig. 3C). In contrast, leg trajectory orientations, captured by the angle of PC1, showed that foreleg and hindleg trajectories are more aligned with the primary body axis in forward compared to backward locomotion, with almost no difference in the middle legs (Fig. 3D). To measure step dynamics in the leg-body coordinate system (Fig. 3G), we measured the angle between the vector connecting the base to the tip of each leg and the primary body axis (Fig. 3E-F). This allowed us to compare the angular span of each leg in forward and backward locomotion (Fig. 3H; see Methods), revealing that in backward locomotion, the middle legs are oriented more anteriorly, and that the angular sector they cover is significantly smaller. These striking differences in leg kinematics between forward and backward locomotion suggest that their speeds were adjusted during evolution by fine-tuning the intricate interactions between position and angular span of the legs. More specifically, the lower speed in the backward gait can be attributed to a smaller step stride in the middle legs, and a reduced side motion resulting from the orientation of the forelegs and hindlegs.

### Two ways of moving backwards: mole crickets have evolved a distinct gait for fast reversal

In response to an aversive stimulus from the front (e.g., a mild tap to the insect’s head; see Methods), the mole crickets adopt a unique backward gait. This recently described leg coordination pattern, termed the backward bound [25], has not been observed in any other insect to date. It comprises an alternation between the middle and hind legs with each pair of left-right legs moving in-phase (Fig. 4A, D). The movement pattern of the front pair of legs is less consistent and robust (Fig. 4D).

**Figure 4.**
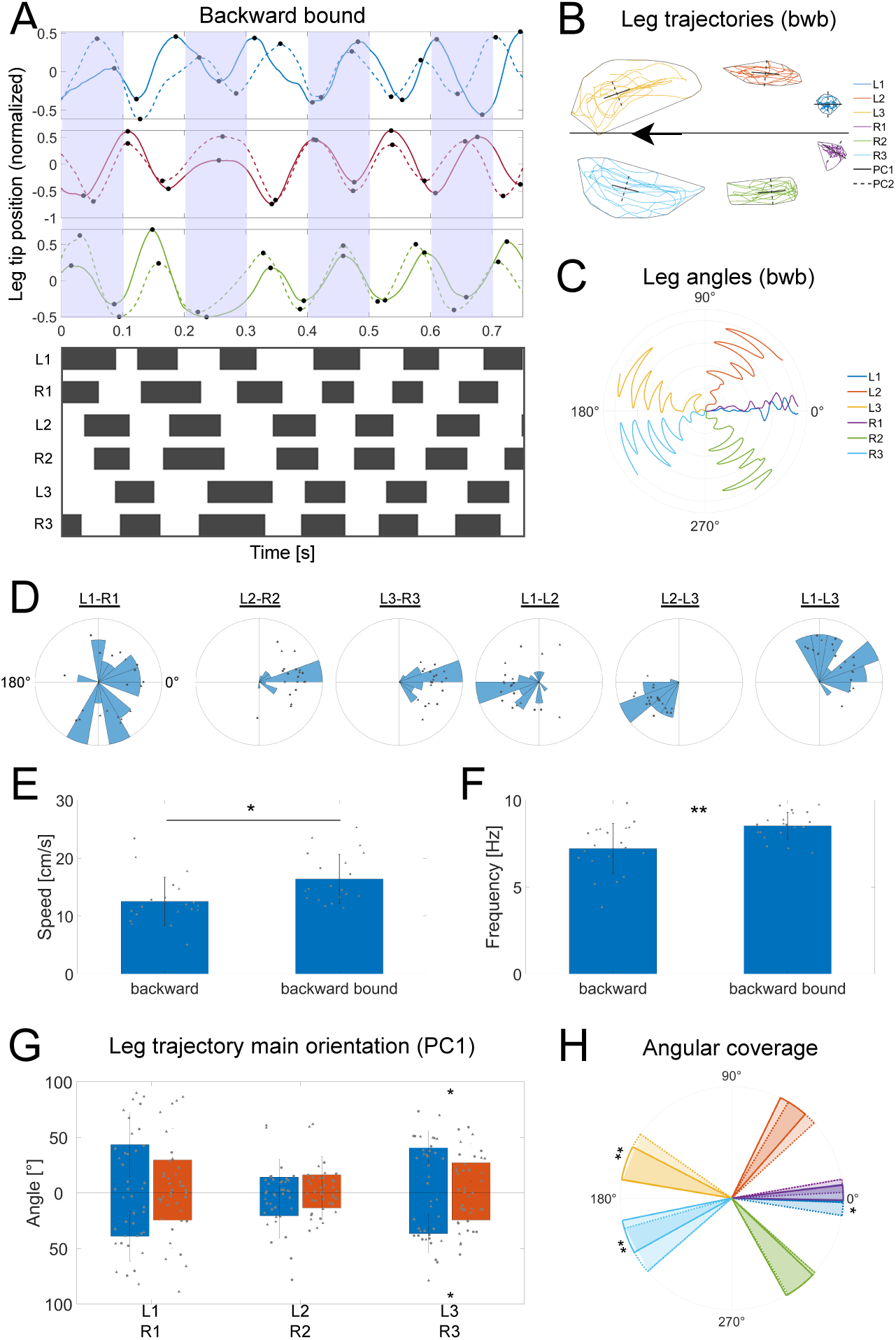
The backward-bound gait. A. Example of leg tip distances over time, *d_i_*(*t*) (see Fig. 2A), in backward-bound, showing the forelegs (top, blue), middle legs (middle, red), and hindlegs (bottom, green), and a corresponding swing-stance diagram (bottom, black bars show stances; white bars show swings; see Methods). B. Example of leg trajectories in the backward-bound (see Methods). The gray contour lines show the minimal convex contour of each leg trajectory. The black solid and dashed lines show the primary (PC1) and secondary (PC2) directions of each leg trajectory, respectively (see Methods). C. An example of leg angles in backward-bound (see Fig. 3G and Methods). Time is shown in the radial coordinate. D. Histograms of inter-leg phases (see Methods). Dots show mean inter-leg phases for single clips. E-F. Whole-body speed (E) and stepping frequency (F). G. Leg trajectory primary orientation (see Methods). H. The mean angular sector covered by each leg in backward (solid lines) and backward-bound (dotted lines) locomotion. The circular sectors show the average standard deviation of leg angles across all animals, and their center shows the average leg orientation (black solid lines for forward and black dashed lines for backward). Statistics were calculated using the two-sample t-test. \**p <* 0.05, \*\**p <* 0.005. Bars show the mean value across animals; error bars show the standard deviation. In H, statistics for the mean angles were calculated using Kuiper’s test. Stars show statistically significant difference between the standard deviations of leg angles. n=20 clips of different animals per gait, comprising at least 8 females (circles) and 8 males (triangles).

Given the use of two distinct gaits for moving backward (either via an alternating-tripod pattern or bounding), we asked whether one has an advantage over the other in terms of whole-body speed. To answer this, we extracted the same spatio-temporal features as above from backward-bound sequences (Fig. 4A-D). We found that, on average, whole-body speed in the backward-bound gait is higher than in the backward double-tripod gait (Fig. 4E), and that this increased speed can be at least partially explained by an increased stepping frequency (Fig. 4F). To determine which leg kinematic properties may further contribute to this difference in speed, we compared the leg trajectories and found that trajectory orientation (PC1) of the hind legs was more aligned with the primary body axis in the backward-bound gait (Fig. 4G). This suggests that the hind-legs might contribute to the ability of the animal to generate more thrust in the backward-bound gait by stabilizing the body and reducing side motion. Moreover, despite the higher stepping frequency (and thus shorter period) in the backward-bound gait, the hindlegs were also found to cover a larger angular sector compared to the backward double-tripod gait (Fig. 4H). Overall, we found that mole crickets move backwards faster in the bound gait compared to the alternating-tripod gait, and that this may be explained by a higher stepping frequency and larger step strides with less side motion of the hind legs. The left-right symmetry in the bound gait might have a stabilizing effect, allowing the animal to push its body backward faster and with less side motion. These findings suggest that the backward-bound gait has evolved and been optimized as a fast escape response. While the slower backward gait (via the alternating-tripod pattern), may be beneficial for navigating and exploring narrow burrows over longer time scales, where turning is either not possible or is energetically costly.

### Modeling the change in angular position and amplitude of middle- and hind-legs is sufficient to reproduce the speed difference between gaits

We have shown that mole crickets employ three distinct walking gaits to navigate their underground tunnels, including one forward gait and two backward gaits. We quantified the spatio-temporal properties of their leg kinematics and revealed differences across these gaits (Figs. 2-4). We also identified differences in whole-body speed between forward and backward double-tripod locomotion, as well as between the two backward gaits. These findings provide insights into their functional roles and the environmental constraints that have shaped them throughout evolution.

However, these data did not allow us to establish a causal link between specific leg kinematics and the ability of the animal to move faster or slower, as other factors may also be involved. Computational models provide a powerful tool for capturing biological phenomena and testing hypotheses that cannot be evaluated in vivo. We combined biomechanical modeling and reinforcement learning to model mole cricket biomechanics and simulate the observed gaits in order to investigate form-function relationships (Fig. 5A-B; see Methods). Reinforcement learning, which is commonly used to solve complex, multi-objective optimization problems [26], was applied here to optimize the state-dependent sequence of joint torques required to replicate the gaits observed in vivo within a physical simulation environment (MuJoCo) [27]. By modifying specific leg kinematics in the model, we were able to predict their contribution to the animal’s ability to generate thrust and propose a causal link between body morphology, leg kinematics, and thrust generation.

**Figure 5.**
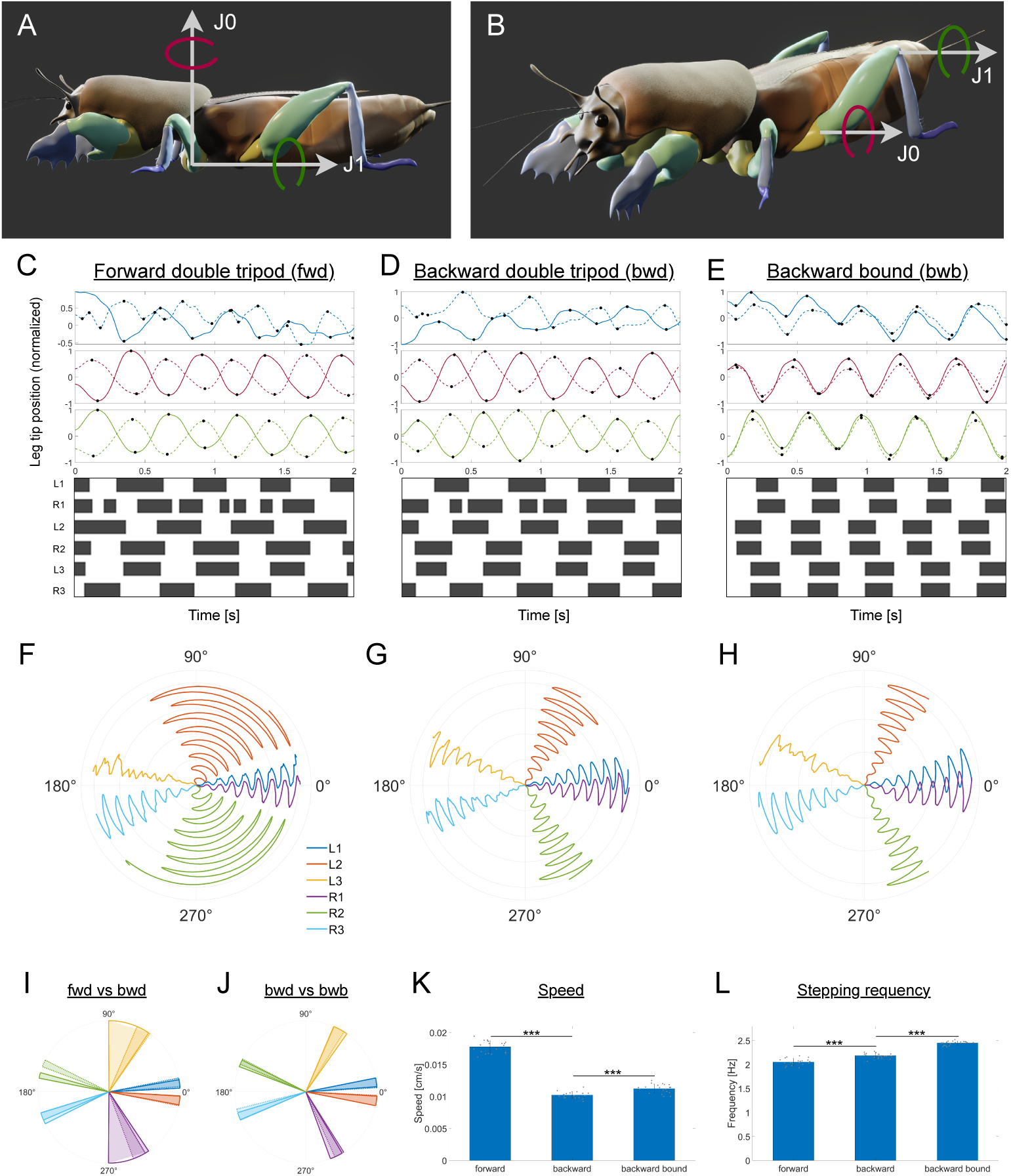
Mechanical modeling and simulation of stepping gaits in the mole cricket. A-B. A 3D model of the mole cricket (see Methods) showing leg segmentation and the joints for which torque was optimized to capture the stepping gaits observed in vivo (Figs. 2-4). C-E. Example of leg tip distances (top) and corresponding swing-stance diagrams (bottom) of forward (D), backward (E), and backward-bound (F) gaits (see Figs. 2-4 and Methods). F-H. Example of leg angles corresponding to C-E, respectively. I-J. Mean and standard deviation of simulated leg angles and comparison between the forward (solid lines) and backward (dotted lines) double-tripod gait (J) and between the backward double-tripod gait (solid lines) and the backward-bound gait (dotted lines) gaits (K). K-L. Whole-body speed (L) and stepping frequency (M) of the simulated gaits. Leg angles and stepping frequency were optimized to match the experimental results (cf. Fig. 3H and Fig. 4H), while speed was not. n=20 simulations per gait. Statistics were calculated using the two-sample t-test. \*\*\**p <* 0.0005. Bars show the mean value across simulations; error bars show the standard deviation.

We used the experimental data to constrain the inter-leg phases, as well as the angle and angular span of joints in the middle and hind legs in our model (Fig. 5C-H). To compare simulation and experimental statistics, we simulated each gait 20 times, initiating the body configuration (i.e., joint angles) randomly within certain error bounds in each simulation (see Methods). However, it should be noted that the variation across individual animals was not modeled here and is beyond the scope of the present work. By applying the same computations and extracting the same kinematic features, we were able to validate our simulations against the experimental data (Fig. 5I-J). Additionally, parameters that could not be measured in vivo, such as joint torques and contact forces, were extracted from the simulations where possible.

To estimate thrust generation, we measured whole-body speed and found that, consistent with our experimental results, forward speed was approximately double that of backward speed, while stepping frequencies were similar (Fig. 5K-L and see Fig. 2D-E). A slight increase in the stepping frequency of the backward-bound gait compared to the backward double-tripod gait was insufficient to generate a faster bound gait. Further observations from a side view revealed that, during the bound gait, the animal raises its body higher [25]. Incorporating this feature was sufficient to generate a faster bound gait (Fig. 5K-L and see Fig. 4E-F).

Importantly, while stepping frequency was set as an objective in the optimization, speed was not. Adjusting the angular position and amplitude of the middle and hind legs was sufficient to capture the speed trends across gaits (in addition to the inter-leg phases, which differ between the two alternating-tripod gaits and the bound gait). By making only these specific modifications, we were able to isolate their effect on locomotion and posit a causal link to the animal’s ability to generate thrust.

To further verify this causal link, we next asked whether the observed speed differences could be eliminated by generating a forward-like backward gait, and a backward-like bound gait, through adjustments to the angular position and amplitude of the middle and hind legs. Indeed, making these adjustments was sufficient to eliminate the speed difference between forward and backward locomotion (Fig. 6A-B), as well as to make the bound gait slower rather than faster than the backward gait (Fig. 6C-D).

**Figure 6.**
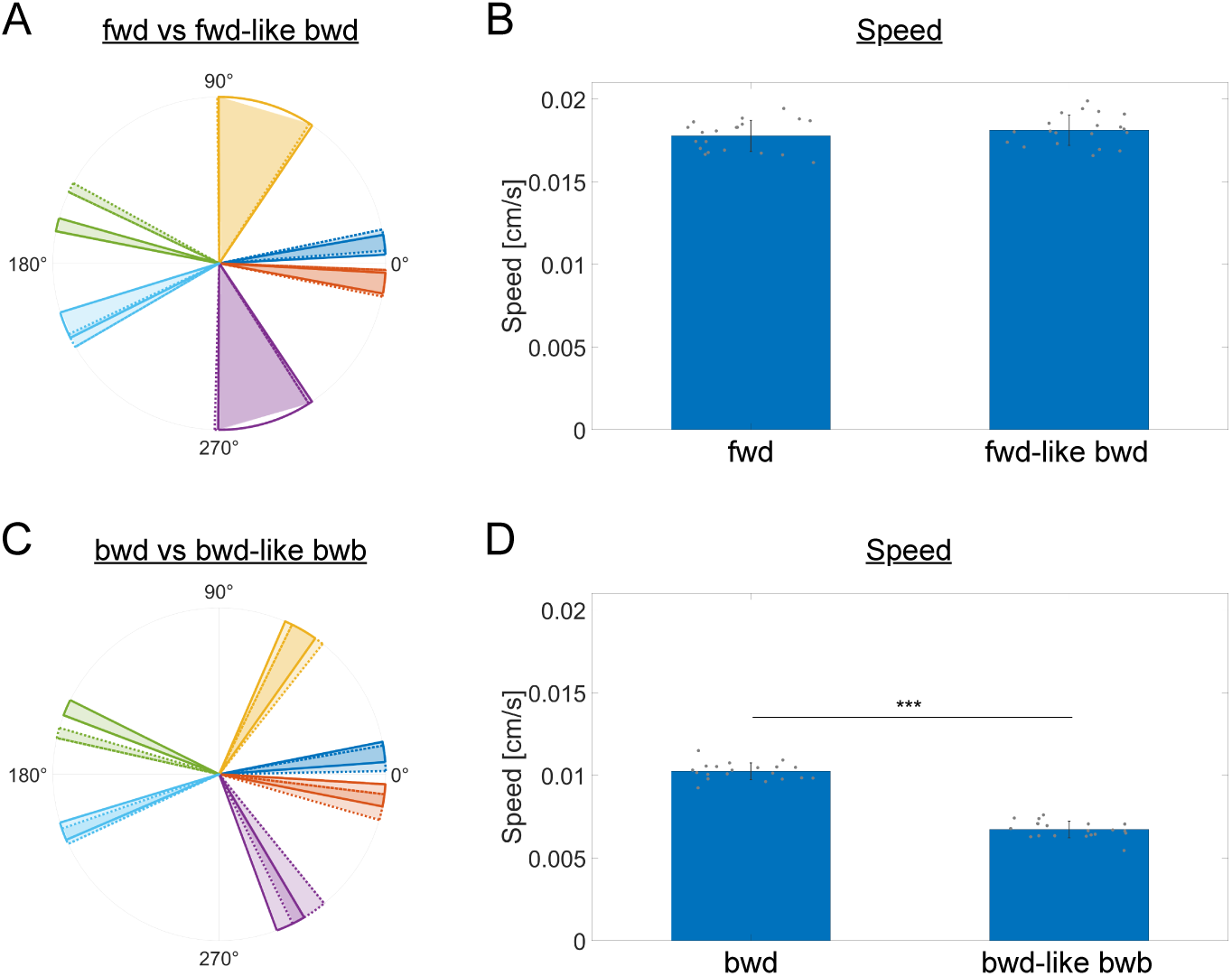
Generating modified gaits to verify model predictions regarding the factors contributing to the mole cricket’s ability to generate thrust. A. Leg angular coverage compared between simulated forward (solid lines) and forward-like backward (dotted lines) gaits, in which the angular position and range of the middle and hind legs were adjusted to match those in forward locomotion. B. Whole-body speed of the forward-like backward gait compared to the forward gait. C. Leg angular coverage compared between simulated backward (solid lines) and backward-like bound (dotted lines) gaits, in which the angular position and range of the middle and hind legs were adjusted to match those in forward locomotion. D. Whole body speed of the backward-like bound gait compared to the backward gait. n=20 simulations per gait. Statistics were calculated using the two-sample t-test. \*\*\**p <* 0.0005. Bars show the mean value across simulations; error bars show the standard deviation.

Overall, by capturing the observed gaits in a controlled physical simulation environment, we were able to isolate the effect of specific leg kinematic properties on the animal’s ability to generate thrust. Furthermore, in addition to capturing the observed gaits in the simulation, we generated modified gaits and showed that eliminating specific leg kinematic differences between gaits (apart from inter-leg phases) is sufficient to eliminate their speed differences.

## 3 Discussion

The study of animal locomotion is of great importance in several fields of science, including ecology, motor control, navigation, decision-making, biomechanics, and robotics. While all insects share structural constraints such as six legs and body segmentation, they represent one of the most diverse groups of animals in terms of their body morphology, locomotion, and biomechanics [1, 28]. Studying this group of animals, which provides a common basis for comparison while exhibiting cross-species diversity due to evolutionary adaptations, offers the potential to uncover universal principles of form-function relationships in locomotion across various environments and contexts, as well as their underlying neural and muscular control mechanisms.

A major challenge in uncovering such universal principles is that of the need to also study the more elusive species that inhabit extreme and less accessible environments. Such species often develop unique adaptations to their habitats, manifested at all levels, from behavior, to body structure, to the neuromuscular control mechanisms that drive their movement. The mole cricket is one such fascinating example. It is a subterranean insect that specializes in digging and navigating a complex network of subterranean tunnels. For this, it has evolved specific adaptations, including a pair of fossorial legs, a strong exoskeleton, and elongated sensory appendages at either end of its body (antennae at the front and cerci at the rear). However, due to its cryptic lifestyle, little is known about the mole cricket to date.

In this work, we studied the multiple locomotion gaits of the mole cricket and compared their kinematic properties in the context of the animal’s unique morphological adaptations. We identified three major modes of locomotion: forward and backward gaits via an alternating tripod pattern, and a backward-bound gait, in which each left-right pair of legs moves in-phase [25].

Analysis of the two double-tripod gaits suggests that the forward gait is faster (despite the similar stepping frequencies) due to a larger stride length in the middle legs, as well as the orientation of the trajectories of the forelegs and hindlegs, which are more aligned with the direction of motion compared to the backward gait. Given that the forward double-tripod gait is much more prevalent in insects, these findings suggest that the leg kinematics of the backward double-tripod gait were adapted to optimize it for its purpose.

A lower speed of backward locomotion may be advantageous in some scenarios, such as for careful sensory integration and decision-making during backward navigation, or exploration in environments where turning is difficult or impossible. The shorter stride length in the middle legs and the increased lateral motion of the forelegs and hindlegs in the backward gait might reflect some morphological constraints. However, they may have also evolved to support the function of this gait. For example, lateral motion may be useful for detecting crossroads while navigating backwards in a complex network of tunnels.

To further examine the link between leg kinematics and whole-body speed, we combined mechanical modeling and reinforcement learning to simulate the observed gaits. We used experimental data to constrain our simulations (matching inter-leg phases, leg angles, and speed and frequency trends). After verifying the simulated models, we generated modified gaits to isolate the contribution of specific leg kinematic properties to whole-body speed. By generating a forward-like backward gait, we demonstrated that matching the stride length (measured by leg angles) in the middle legs in the backward gait to that in the forward gait was sufficient to eliminate their speed difference.

Next, following the identification of a six-legged backward-bound gait that has not been observed in any other insect [25], we asked why mole crickets have evolved two distinct backward gaits. A comparison of the two gaits revealed that both speed and frequency were slightly higher in the bound gait. Furthermore, bound sequences typically included fewer step cycles. Taken together, this suggests that the backward-bound gait has evolved as a transient, fast escape mechanism for distancing the animal from an immediate threat.

Given these findings, we then asked which leg kinematic properties facilitate the increased speed in the backward-bound gait compared to the backward double-tripod gait. Importantly, the higher stepping frequency in the bound gait is a major potential contributor to the increased speed at the neural control level. In addition, the in-phase motion of contralateral leg pairs on its own may have a balancing effect, enabling the animal to generate more thrust compared to the double-tripod gaits. However, simulation of the bound gait did not result in higher speed (data not shown), suggesting that stepping frequency and inter-leg phases in the bound gait alone are insufficient to account for its increased speed.

Finally, we used observations from a side view, which revealed that during the backward-bound gait the animal raises its body higher compared to the double-tripod gaits. Adding this constraint to the bound gait simulation was sufficient to produce a faster gait. Conversely, removing this property such that body lifting became similar in both backward gaits, resulted in a bound gait that was slower than the backward double-tripod gait.

In this work we identified and characterized the different walking gaits in the mole cricket. Quantitative analysis of these gaits revealed striking differences in speed and stepping frequency, as well as in leg kinematics, manifested through either temporal or spatial properties. Although this analysis included both male and female mole crickets, no significant differences were found between the two sexes. In order to answer form-function questions that cannot be addressed using in vivo experiments alone, we captured the observed gaits in a physical simulation environment using reinforcement learning and generated modified gaits in order to pinpoint the contribution of specific leg kinematic properties to the ability of the animal to generate thrust. Importantly, modeling predictions should generally be interpreted with caution, as the model and simulations are only approximations of the animal and its gaits. In particular, in our framework, the joint mechanics and actuation were modeled using a simple spring-damper system, which does not capture the full complexity of biological joints and muscles. Additional limitations of the model include simplified body geometry (e.g., small protrusions and hairs are ignored), mechanics (e.g., the body lacks flexible parts), and forces at the body-environment interface (e.g., friction and adhesion).

## 4 Methods

### Animal maintenance and experimental procedure

Mole crickets (*Gryllotalpa sp.*) were collected at different life stages from the field in the countryside around Tel Aviv, Israel. They were maintained at room temperature (20–25^◦^C), each in a separate 750 ml glass jar filled with autoclaved plant soil, and fed on flour beetle larvae, grass roots, and slices of carrot. The soil in each jar was moisturized by sprinkling 100 ml of water every three days. Animals (adults only) were acclimated to the laboratory conditions for at least 1 month before onset of the experiments. The imaging setup consisted of a tunnel-like compartment (30L x 2.5W x 10H cm) made of foam-board, with an acclimation chamber (10L x 10W x 10H cm) made of 4-mm-thick acrylic, polymethyl methacrylate board (PMMA, Perspex) attached at each end (Fig. 1C). A millimeter paper was glued to the base of the tunnel in order to extract a scaling factor to convert from pixels to real length units (Fig. 1C). The width of the mole cricket’s tunnel in nature is 2-3 times its thoracic width [29]. For this study, we set the tunnel width to 2.5 cm, which is approximately 2.5 times the average body width of an adult mole cricket, as measured at the pronotum’s broadest part, (10.06 ± 0.83mm for males and 9.9 ± 0.09mm for females) [17]. Recording was carried out from above (∼ 20 cm above the base of the container) using a high-speed camera (MIKROTRON motion-Blitz cube4MGE-CM4, Germany) equipped with lens for focusing (Vital Vision Technology Pte Ltd, Singapore), at 500 fps, at a resolution of 1280 x 1064 pixels, and a temperature of 24 ± 1^◦^C. The scene was illuminated by a dual 20W goose-neck fiber optic lamp, with an additional LED bulb, (IKEA Jansjo desk lamp - flexible goose-neck lamp) to prevent shading. The bulbs were covered with red cellophane to reduce both glare and stress to the insect. Adult male and female mole crickets were introduced individually into one of the acclimation chambers, and their locomotion was recorded for up to 10 minutes while they were in the tunnel section. Motion was triggered by tapping on the animal’s head from the front or the cerci from behind, using a sponge attached to a wooden stick. The force exerted by this stimulus was not sufficient to move the animal’s body on its own. For each gait, 20 videos of different animals were recorded, incorporating at least 8 males and 8 females. Videos were considered valid if they contained at least three step cycles.

### Video analysis and extraction of locomotion parameters

To extract kinematic parameters from the recorded data, key points on the animal’s body were tracked in all videos, comprising leg tips (tarsus), leg bases (thorax-coxa junction), and the thorax-head (TH) and thorax-abdomen (TA) junctions (Fig. 2A). A total of 1,512 frames were selected from 24 videos for key point labeling to train a ResNet50 model. Labeled frames were selected from videos of different gaits, body poses, and animals. The trained model was then applied to the entire video dataset to detect these points in all frames (n=15,878 in 60 clips). Where needed, key points were manually corrected (n=275 frames, 1.7%), mostly in cases of self-occlusion and soil particles attached to the body. Annotation, training, and detection were carried out using DeepLabCut (DLC) [30].

The detected points were then used to define a body coordinate system and quantify leg kinematics. The main body axis was defined as the vector connecting TA to TH (see yellow arrow in Fig. 2A). The secondary body axis is perpendicular to the main body axis, with its origin at the TA point (see yellow line in Fig. 2A). Whole-body speed was defined as the change in the position of the TA point per unit time, and the TH point was used to determine the sign of the speed (i.e., forward or backward). This was then averaged across all frames of a single clip, and then averaged again across clips of the same gait. Leg tip position of leg *i*, *d_i_*(*t*), was defined as the shortest distance of the leg tip from the secondary axis. The peaks in the graph of *d_i_*(*t*) were used to compute step cycle period and stepping frequency. In each clip, this was first computed for each leg, then averaged across all legs, and then averaged across all clips of the same gait. Using the maxima and minima in the graph of *d_i_*(*t*), forward stances were defined as the time ranges from the anterior-most to the posterior-most leg tip positions, and swings were defined as the remaining time intervals. Backward stances were defined in the opposite way (i.e., from the posterior-most to the anterior-most leg tip positions).

The inter-leg phase between legs *i* and *j* was calculated by computing the cross-correlation between *d_i_*(*t*) and *d _j_*(*t*), and defined as the phase corresponding to the highest peak (in absolute value) in the correlation-phase graph. The trajectory of a leg was defined as its tip position over time, in which the coordinate in each frame was translated such that the TA point is at the origin, and rotated such that the main body axis is aligned with the positive x-axis. To estimate the area covered by each leg trajectory, the minimal convex contour was computed (see gray polygons in Fig. 3A-B). PCA was used to decompose each polygon into its primary and secondary components. The primary component was used to capture the main orientation of the trajectory and approximate step stride (defined as the maximum distance along the first component within the polygon). Leg angle was defined as the angle between the vector connecting the leg base to its tip, and the main body axis (see Fig. 3G). To obtain the average leg angle, the circular mean was first used to average leg angles over time and then across videos. Similarly, the circular standard deviation was used to obtain the standard deviation of leg angles over time, and then averaged across videos using a regular mean.

### Biomechanical modeling and simulation of walking gaits

A 3D geometrical model of the mole cricket was designed using Blender 3.6 [31]. The model is bilaterally symmetrical, and segmented such that the head, thorax, abdomen, and legs were made of separate meshes. Each leg was further segmented into the coxa, femur, tibia, and tarsus. Next, all body parts were exported from Blender in .stl format and imported into the MuJoCo physical engine (Multi-Joint dynamics with Contact) [27]. In MuJoCo, the body parts were connected via hinge joints (1D spring-damper joints), and each joint was associated with a motor enabling torque to be applied to the joint. Specifically, two orthogonal joints were placed between the coxa and femur, one between the femur and tibia and one between the tibia and tarsus (Fig. 5A-B). The rotation origin and axis of each joint were chosen such that they approximately matched those of the animal as derived from our observations. Finally, mechanical parameters such as body mass, joint damping, and friction were set.

To find a state-dependent sequence of joint torques that captures a certain gait, we used the PPO (Proximal Policy Optimization) algorithm, which is a reinforcement learning (RL) agent widely used for solving motion-planning problems with a high degree of freedom [32]. For the policy (actor) and value (critic) neural networks, we used MLP (multilayer perceptron) networks with two hidden layers, each with 128 neurons. tanh() was used as the activation function in both networks. The reward (objective) function consisted of three main terms: a constant step (survival) reward; a high-level forward reward to motivate the agent to progress in the positive y-direction; and a low-level imitation reward to impose a soft constraint on inter- and intra-leg phases with a target frequency. For the reference imitation data of the different gaits, we used a sine function to impose a constraint on the joint angles. During training, the reference phase of the step cycle was continuously adjusted by taking a weighted average of the reference phase with the phase that best matched the current body configuration. This was crucial to account for physical interactions with the environment and noise, as the agent would otherwise lose synchronization with the reference phase. Finally, a termination condition was set such that an episode would terminate if the body configuration crossed thresholds of the reference inter- and intra-leg phases. This technique prevents divergence and is widely used in motion-planning problems to narrow the exploration space to relevant solutions [26]. The observation space (i.e., the information available to the RL agent, based upon which it makes decisions) included joint angle and joint angle velocities.

Training of the PPO agent was performed using a time step of 0.01s for the optimization and 0.001s for the physics. The episode duration was set to twice the period of the target gait. The program was run on a Windows 11 machine with a NVIDIA RTX6000 GPU, which was used to train 16 RL environments in parallel. Each training session was run for a maximum of 20 million steps, which took approximately 4 hours. The full code is freely available at https://github.com/Omer1Yuval1/insectRL.

